# Prioritizing Management for Cumulative Impacts

**DOI:** 10.1101/2021.10.07.463502

**Authors:** Gerald G. Singh, Jonathan Rhodes, Eve McDonald-Madden, Hugh P. Possingham, Edd Hammill, Cathryn Clarke Murray, Megan Mach, Rebecca Martone, Benjamin Halpern, Terre Satterfield, Kai M. A. Chan

## Abstract

Determining where environmental management is best applied, either through regulating single sectors of human activities or across sectors, is complicated by interactions between human impacts and the environment. In this article, we show how an explicit representation of human-environment interactions can help, via “impact networks” including activities (e.g. shipping), stressors (e.g. ship strikes), species (e.g. humpback whales) or ecosystem services (e.g. marine recreation). Impact networks can enable the identification of “leverage nodes”, which, if present, can direct managers to the activities and stressors crucial for reducing risk to important ecosystem components. Exploring an impact network for a coastal ecosystem in British Columbia, Canada, we seek to identify these leverage nodes using a new approach employing Bayesian Belief Networks of risks to ecosystems. In so doing, we address three key questions: (1) Do leverage nodes exist? (2) Do management plans for species correctly identify leverage nodes? (3) Does the management of leverage nodes for certain species realize benefits for other species and ecosystem services? We show that there are several leverage nodes across all species investigated, and show that preconceptions about the regulation of risk to species can misidentify leverage nodes, potentially leading to ineffective management. Notably, we show that managing fisheries does not reduce overall risk to herring whereas managing diverse cumulative impacts including nutrient runoff, oil spills, and marine debris can reduce risk to herring, additional species, and related ecosystem services. Thus, by targeting leverage nodes, managers can efficiently mitigate risks for whole communities, ecosystems, and ecosystem services.

## Introduction

Ecosystems are made up of interacting components and face multiple anthropogenic stressors (1, 2). Effective and efficient management thus requires understanding the processes by which cumulative impacts occur, and the ability to locate key intervention points (3–5). For example, globally marine ecosystems are at risk from anthropogenic activities occurring on land and at sea (1), and effectively managing these ecosystems will require managing these diverse activities across a range of sectors, including fisheries, aquaculture, and marine construction (6, 7). Few management plans are designed to regulate the human activities that cause impacts across economic and industry sectors, but determining when and where sectoral versus integrated management is needed has not often been shown empirically (7, 8). We propose analyzing networks – representing human activities and pressures as nodes that drive impact and environmental components as nodes that receive impact – to help identify and prioritize key activities and stressors for management.

Environmental managers routinely, and often implicitly, assume that certain nodes in a directional network linking human activities with environmental impacts disproportionately regulate risk to ecosystem components (e.g., species, habitats), offering the greatest leverage for management action. But do such “leverage nodes” exist, or is risk to ecosystem components distributed relatively evenly across a network of causal pathways? If such leverage nodes exist, then management can benefit from prioritization schemes that identify and target them. If instead they are rare or do not exist, then management should regulate risk across an impact network.

Cumulative impacts to ecosystems from multiple activities have been analyzed using networks based on causal pathways of impact, including identifying nodes (e.g., pressures) with the most connections to other nodes (e.g., species), and to identify mechanistically similar pathways (3, 9), but they do not account for the magnitude of impacts. More recently, networks have been constructed with associated impact weights, characterizing cumulative impacts in marine systems (10). However, impact weights have only been calculated to identify the nodes that contribute impact across an ecosystem, and as of yet no method has been developed to determined what nodes are most important in regulating impact to particular components of the environment that are priorities.

Here we use Bayesian Belief Networks (BBNs) to determine nodes that regulate disproportionate risk to valued ecosystem components (“leverage nodes”). We analyze leverage nodes for coastal British Columbia, where a risk analysis protocol based on causal pathways was developed for seventeen marine species contributing to fifteen ecosystem services (11). We create a network of these causal pathways to track the contribution of risk through different pathways and calculate summed total risk. The causal pathways link human activities (or global stressors) to specific pressures to species to ecosystem services, and we use this structure to determine leverage nodes – nodes that contribute over twice the average total change in risk probabilities to guarantee target species or species groups are categorized as low risk – using BBNs (see Methods for more detail). Using the management of herring (*Clupea pallasii*) as a case study, we compare the leverage nodes revealed from our analysis for herring (revealed leverage nodes) to the targets of management action for a current herring management plan (assumed leverage nodes). Herring is the target of an important commercial, recreational, and cultural fishery in BC (27).

To determine if assumed or revealed leverage nodes provide better management outcomes, we utilize a BBN to determine risk outcomes for herring given management of (*i*) assumed leverage nodes and (*ii*) revealed leverage nodes. The assumed leverage nodes were determined by identifying the management focus of the herring Integrated Fisheries Management Plan (IFMP) in British Columbia (12). We also use this comparison of management for revealed and assumed leverage nodes for herring to determine if management of revealed leverage nodes reduces risk for other species and associated ecosystem services compared to management plans based on assumed leverage nodes. Further, we evaluate the management outcomes of managing for leverage nodes across all species at risk (rather than specific species) to determine if management based on assumed or revealed leverage nodes across a collection of species perform better. Finally, because regulating for leverage nodes is dependent on learning which nodes have high leverage for regulating risk, we explore how leverage nodes can be used in an adaptive management framework – where updated knowledge on leverage nodes at a second time step may affect management effectiveness. In contrast, the IFMP is not built on an adaptive management framework. We use the risk assessment protocol to establish three scenarios for a second time step. The first scenario follows the results of managing assumed leverage nodes under the IFMP, and continues managing these assumed nodes in the second time step. The second scenario follows the results of the revealed leverage nodes for herring and determines new revealed leverage nodes to manage for the second time step. The final scenario follows the results of the revealed leverage nodes of multiple species at risk and determines new revealed leverage nodes for the second time step. This ability to analyze scenarios and evaluate different risk probabilities at simulated sequential time steps expands and improves on previous approaches to analyze networks of cumulative impacts (3, 9). Our analysis explores whether managing assumed or revealed leverage nodes leads to more effective management, and we show that by determining performance of different management plans managers can determine if single-sector or multi-sector management is more effective for a given management challenge.

## Results

### Determining Revealed Leverage Nodes in Individual and Groups of Species

In total there are 86 nodes in the network as identified through the risk assessment (11). The number of activity and pressure nodes associated with individual species ranges from 35-48. Leverage nodes (contributing over twice the average risk to species) were found for each of the 11 species we investigated. The number of leverage nodes per species varied from 1 to 11 (comprising 3-23% of total nodes contributing to risk to each species), with humpback whales having the fewest and seagrasses having the most revealed leverage nodes (Table S3). The mean number (± standard error) of leverage nodes per species in our sample is 6.91 (±1.02). In every case, leverage nodes are found across sectors, and in 64% of species we found leverage nodes across all four sectors that generate impact (fisheries, other sea-based activities, land-based activities, and long term stressors). The SAR in our sample (humpback whale, Cassin’s auklet, Steller sea lion) have relatively few leverage nodes (1, 2, and 3 respectively), each of which can be managed by focusing on a subset of sectors.

We determined that most leverage nodes were stressor nodes. We found 18 leverage nodes across the total set of stressor nodes and 1 leverage node within the activity nodes across our sample of species. For example, if a species is primarily affected by two stressors (e.g. harvesting and habitat alteration), and these two stressors are primarily driven by a single activity, such as bottom trawl fisheries, then the activity node for bottom trawl fisheries will be a leverage node. In contrast, leverage nodes can also be found at the stressor stage. For example, if a species is primarily affected by acoustic stress, and there are multiple shipping and other activities contributing to acoustic stress, the node of acoustic stress is a leverage node. Potential management actions on this node could be limiting boat activity where the species occurs, or applying vessel quieting technologies to the vessels that contribute to acoustic noise (13).

In order to manage SAR along with herring at low risk (*SAR Leverage*), there are 5 leverage nodes (10% of the 48 nodes contributing risk to these species). There are fewer leverage nodes in the *SAR Leverage* scenario than the *Herring Leverage* scenario, but the leverage nodes common to both require a greater decrease in risk for the *SAR Leverage* scenario. For example, oil spills and acoustic stress have more weight as leverage nodes in the *SAR Leverage* scenario compared to the *Herring Leverage* scenario. However, in the second time step, the *SAR Leverage* scenario has more leverge nodes than the *Herring Leverage* scenario (11 in *SAR Leverage*, 8 in *Herring Leverage*).

### Comparing Assumed and Revealed Leverage Nodes

The BBN analysis of the *Herring Leverage* and *SAR Leverage* scenarios allowed us to compare management performance of regulating revealed leverage nodes with the assumed leverage nodes within the *IFMP* scenario. We found that the risk level probabilities of 35 activity and 54 stressor nodes in the impact network changed from initial conditions to the *Herring Leverage* scenario. Of these, we find 7 leverage nodes for herring (Table S3): direct capture, acoustic stress, oil spill, change in water flow, marine debris, temperature change, and nutrient input. The identity of leverage nodes is highly dependent on the variation in risk levels available for management to act on, and less dependent on the probabilities a given species is at a given risk level (Table S7, Table S8).

In comparing the revealed leverage nodes of the *Herring Leverage* scenario with the assumed leverage nodes of the *IFMP*, we find little overlap. First, the *IFMP* assumes the leverage nodes to be three drivers (seine fisheries, recreational fisheries, gillnet fisheries), and one stressor (bycatch). Our analysis ranks these assumed leverage nodes as the 10^th^, 13^th^, 14^th^, and 28^th^ most influential nodes for herring, respectively (out of 54 total, Table S4). The *IFMP* effectively reduces direct capture by keeping three types of fisheries low, but does not address many of the other influential nodes, which may be important priorities to ensure management efficacy (Figure S2).

The *Herring Leverage* scenario greatly increases the probability of herring facing low risk compared to initial conditions (~40%). Both scenarios also increase the likelihood of herring facing high risk (~30% and 25%, respectively). In contrast, the probabilities of herring being at low or high risk in the *IFMP* is 15%. Both scenarios also increase the likelihood of herring facing high risk (~30% and 15%, respectively), compared to initial conditions (11% low risk and 11% high risk). In both scenarios there is a reduced probability of medium risk (30%, and 73% respectively) compared to initial conditions (79% medium risk, Figure 1 Time 1).

**Figure 1.**
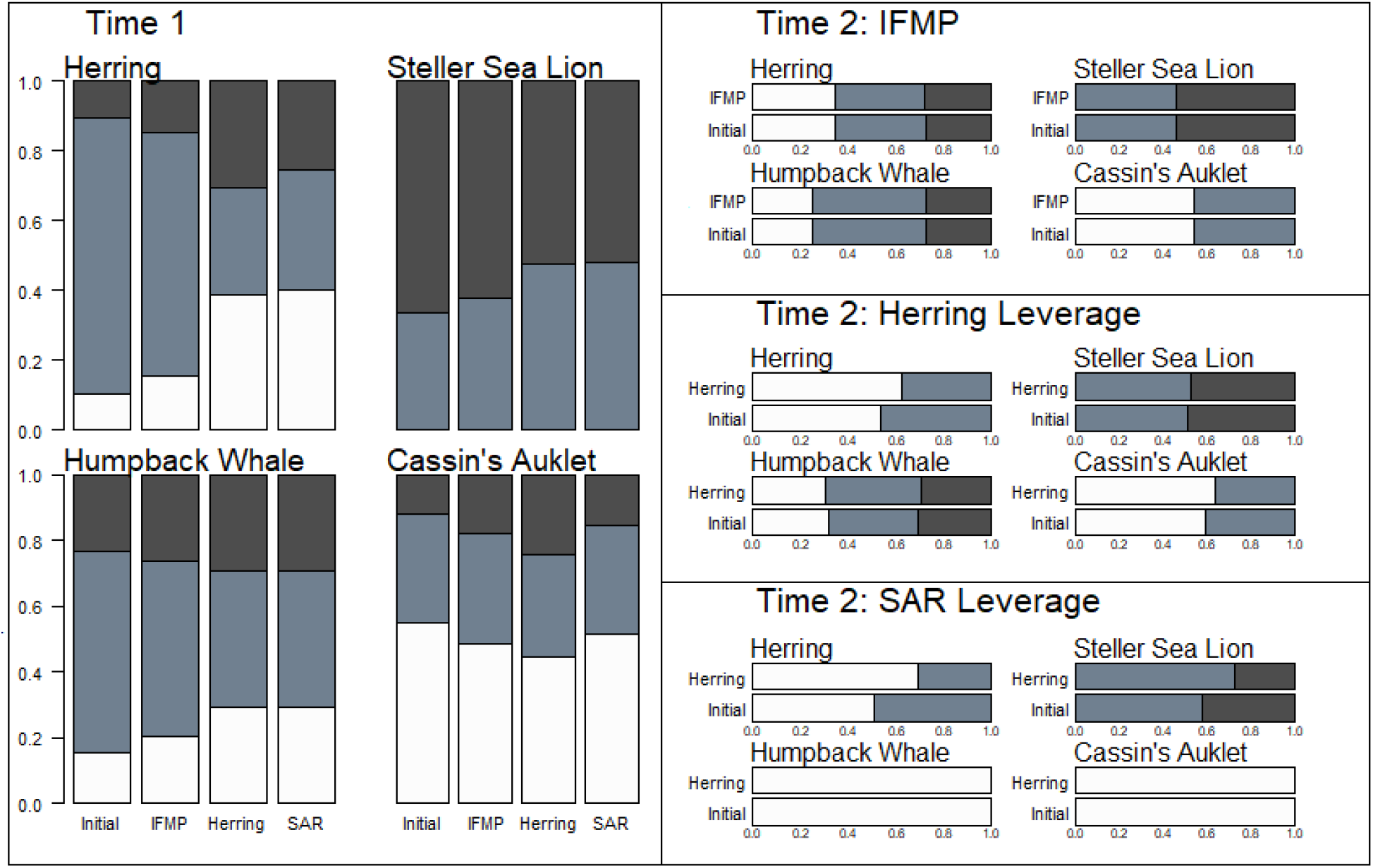
The probabilities of low, medium, and high risk for four species depending on the five different scenarios, at times 1 and 2. White represents low risk, medium grey represents medium risk, and dark grey represents high risk.

The *IFMP* scenario only slightly changes the risk probabilities of other species at risk, and actually leads to an increased probability of high risk for Cassin’s auklet and a reduced probability of low risk compared to initial conditions. The *IFMP* scenario also produces no significantly differences for the number of species or the number of ecosystem services provided at low, medium, and high risk compared to initial conditions. In contrast, the *Herring Leverage* scenario leads to greater changes in the risk probabilities across species and ecosystem service provision. For all species at risk the *Herring Leverage* scenario leads to greater probabilities that the species faces low risk (medium risk for Steller’s sea lion, which only faces medium and high risk) compared to the *IFMP*. However, like *IFMP*, the *Herring Leverage* scenario leads to Cassin’s auklet facing lower probabilities of low risk than initial conditions. It also leads to slightly higher probabilities that these species face high risk than the *IFMP*. The *Herring Leverage* scenario also has significantly more species facing low risk than the *IFMP* though there are no differences in the provision of ecosystem services.

Differences between *IFMP* and *Herring Leverage* are more prominent in the second time step (Figure 1 and 2, Time 2). While total risk across many species is lower at the second time step because the scenario assumes that all the assumed leverage nodes were effectively regulated, the *IFMP* at the second time step produces the same probabilities of species and ecosystem service provision compared to time 2 initial conditions. In contrast, the *Herring Leverage* scenario produces further reductions in risk across species and ecosystem service provision. Similar to the *IFMP*, time 2 for *Herring Leverage* assumes that revealed leverage nodes at time 1 were regulated (so total risk to species is lower at time 2 than at time 1), however unlike the *IFMP* scenario the *Herring Leverage* scenario identifies a new set of leverage nodes at time 2. At time 2 the revealed leverage nodes for *Herring Leverage* are nutrient input, direct capture, marine debris, sedimentation, debris, oil spill, temperature change, and persistent organic pollutants (table S3). Managing these new leverage nodes leads to greater probabilities of low risk across species and ecosystem service provision in the *Herring Leverage* scenario compared to the *IFMP* scenario.

**Figure 2.**
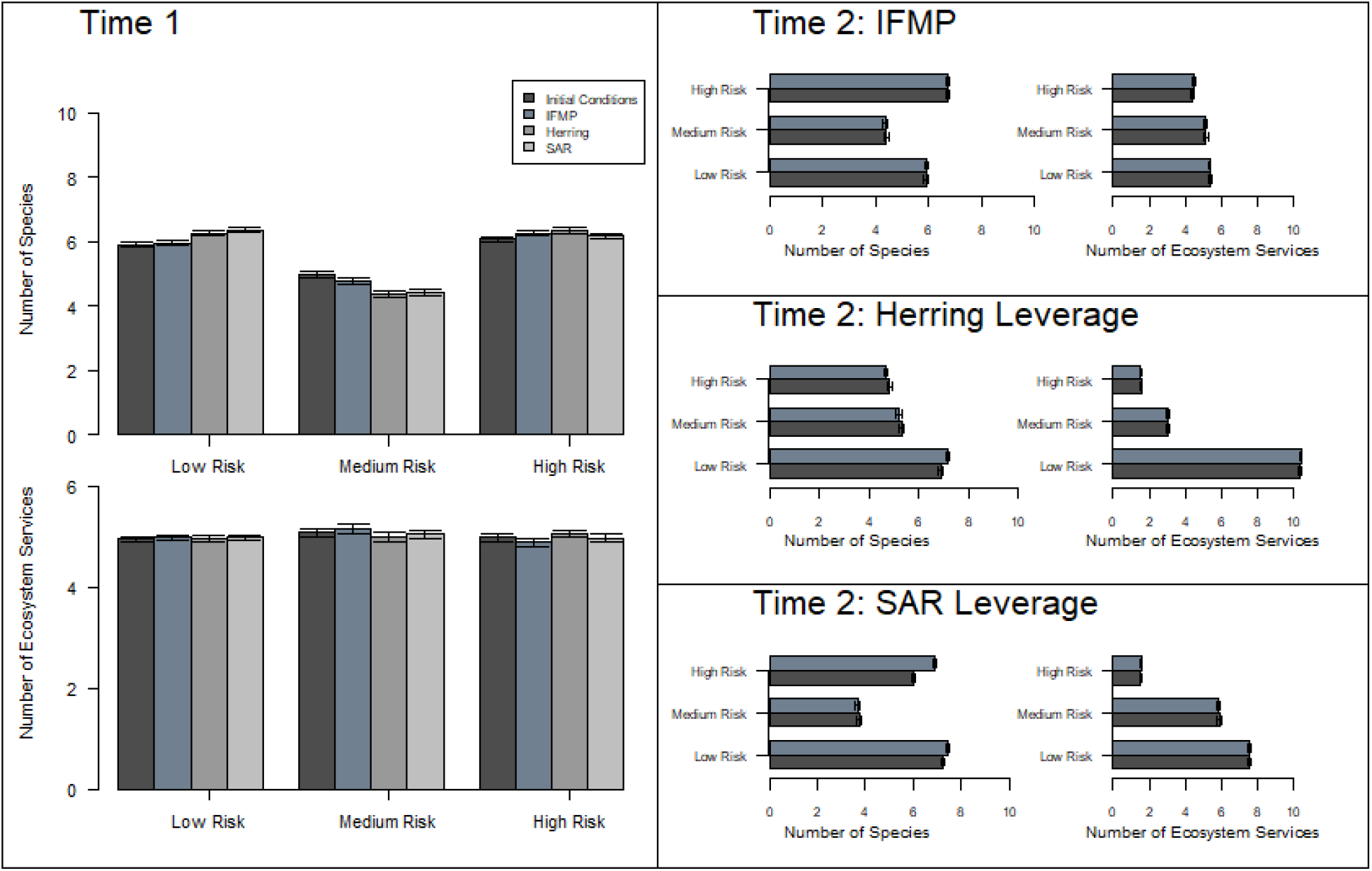
The number of species and ecosystem services predicted to be at low, medium, and high risk from the five different scenarios, at times 1 and 2.

### Comparing Single Species Management versus Multi-Species Management

Under the *SAR Leverage* scenario, herring and the species at risk are more likely to face low risk compared to the *Herring Leverage* scenario. Steller sea lion faces equal risk in these two scenarios. The *SAR Leverage* scenario also leads to more species facing low risk and fewer species facing high risk compared to the *Herring Leverage* scenario, though this difference is marginal. There is no difference in the number of ecosystem services facing different risk levels.

At time step 2 the differences between the *Herring Leverage* and *SAR Leverage* scenarios are more pronounced. At time 2 the revealed leverage nodes are oil spill, marine debris, debris, direct capture, persistent organic pollutants, habitat disturbance, acoustic stress, temperature change, contaminants from large vessels, incidental mortality from large vessels, and sediment (table S3). The probability of herring facing low risk and Steller sea lion facing medium risk is lower under the *SAR Leverage* scenario, and humpback whales and Cassin’s auklet both are guaranteed to face low risk under the *SAR Leverage* scenario, which is not the case under the *Herring Leverage* scenario. The results across all species and ecosystem service provision at time 2 are more nuanced. While *SAR Leverage* has slightly more species facing low risk, it also has more species facing high risk compared to *Herring Leverage*. A greater number of ecosystem services also face low risk in the *Herring Leverage* scenario at time 2 compared to *SAR Leverage*.

## Discussion

For every species and every multi-species scenario (including in a sensitivity analysis, Tables S7 and S8)) we investigated in this coastal BC example, we found nodes (drivers or stressors) that are particularly promising for management (“leverage nodes”). We found between 1 and 11 leverage nodes for species, and did not find that targeting multiple species increases the number of leverage nodes compared to the individual species within the multispecies assemblage. Though there may be cases where managing multiple species has a larger number of leverage nodes, where there are few common prominent activities and stressors that regulate risk among species, management can focus attention on reducing risk from a few activities and stressors. The widespread existence of leverage nodes in our study suggests that management targeted on a few nodes can offer considerable leverage for reducing risk. Encouragingly, we found relatively few leverage nodes for SAR (1-3 nodes), indicating that effectively managing these species may be relatively straightforward. Because species at low population sizes are often vulnerable to a suite of diverse activities and stressors (14), this finding is surprising but may reflect the fact that SAR in our context are species that happen to be affected by a limited number of prominent risk pathways. The existence of leverage nodes supports the approach of many management plans, such that management is focused and not diffuse across the entire impact network (15). However, for every individual species and group of species we examined, we identified at least one leverage node that is cross sectoral, and many of these require management across all four categories of sectors (fisheries, sea based activities, land based activities, and long term stressors). The universality of cross-sectoral leverage nodes challenges the sector-specific approaches employed in many management plans (15).

Our results suggest that prioritizing management based on leverage nodes revealed by impact analysis and BBNs might provide better management outcomes than managing based on assumptions and jurisdictional constraints. When leverage nodes exist at the stressor level, effective management requires managing all prominent activities that drive this stressor or directly mitigating this stressor. Through BBNs and a comparative risk assessment, we show that the *IFMP* could effectively reduce risk from the stressor “direct capture” (the most influential leverage node), but that it does not account for other influential stressors that impact herring such as acoustic stress and oil spill (16, 17). The finding that herring face greater probabilities of both low and high risk under the *IFMP* compared to initial conditions suggests that other drivers of risk can more than compensate for the reduction of risk to herring from direct capture. That is, managing direct capture is not enough to be certain of controlling risk to herring. Considering risk to all assessed species and the provision of ecosystem services, the *IFMP* is indistinguishable from initial conditions, except that there are a higher number of species predicted at high risk and a lower number of species at medium risk. Thus, by focusing only on direct capture and ignoring other stressors, this analysis suggests that the *IFMP* offers little to no benefit for herring or other species (18). Our results show that managing assumed leverage nodes (*IFMP scenario*) do not change risk profiles for any species compared to initial conditions at the second time step, while managing revealed leverage nodes (*Herring leverage* scenario) reduces risk to all species across both time steps. The benefits of determining leverage nodes and managing these revealed leverage nodes as opposed to assumed leverage nodes is more apparent over time. Our results showcase the utility of the method we developed: prioritizing new leverage nodes based on analysis at each time step allows for flexibility and adaptation in management planning, and evaluating multiple management choices across time can be seen as simulating adaptive management (19).

Our analysis suggests that management prioritization based on revealed leverage nodes might have beneficial outcomes, especially when prioritizing multiple commercial species and species at risk. The benefits of targeting revealed leverage nodes is more prominent after multiple time steps, when key leverage points in human activities and stressors are serially acted on and risk is reduced. At the first time step, management of leverages nodes in the *SAR leverage* scenario produces lower risk across species at risk as well as herring compared to the *herring leverage* scenario. There are also more species and ecosystem services at low risk under the *SAR Leverage* scenario. These effects are more pronounced at the second time step, where there are very high probabilities of herring and species at risk facing low risk (and produce greater changes towards low risk compared to initial conditions compared to *Herring leverage* scenario – even for herring), and where there are more species and ecosystem services facing low risk relative to initial conditions compared to the *Herring leverage* scenario. Our results indicate that management for multiple species may be more beneficial than single species management, even for individual species.

More research is needed to fully outline the conditions where multi-species management is preferable to single-species management, but our methods and results highlight some possibilities. By incorporating ecological interactions in the risk assessment, we could address common threats to species through trophic and habitat links (20). For example, all species of concern in the *SAR Leverage* scenario (herring, Steller’s sea lion, Cassin’s auklet, Humpback whale) are dependent on zooplankton directly or indirectly (with one trophic level removed). By focusing across species, the *SAR Leverage* scenario targeted human activities and stressors that contribute risk to this common trophic base (such as contaminants from ships and debris) across time steps as well as target human activities and stressors important to individual species (e.g. direct capture for herring, incidental mortality for Steller’s sea lion). Beyond the focus on multi-species management, our results highlight the importance of taking an ecosystems approach to analysis and management (5, 21). While simply managing more activities and stressors overall may seem intuitively better for ecosystems broadly, our results show that a more nuanced approach is necessary. In the first time step the *SAR Leverage* scenario had fewer leverage nodes than the *Herring Leverage* scenario, while in the second time step it had more, yet in each case the *SAR Leverage* scenario performed better. Determining the correct leverage nodes based on an understanding of the underlying ecology is more important for effective management of an ecosystem, beyond simply managing more nodes.

Though potentially costlier than managing single stressors, managing diverse stressors is more likely to reduce risk to multiple species than managing single, or similar, stressors (8). Traditionally, management for fisheries relies on regulating catch, with little consideration of other impacts on fish populations (21). The focus on selective versus nonselective fishing practices in the fisheries management literature (22) may neglect other important stressors important for fisheries management. All scenarios that include the management of diverse stressors show better risk outcomes for the ecological community than the *IFMP* scenario where functionally similar stressors contributing to catch are managed. We believe two complementary processes contribute to this finding. First, there are key stressors that are mechanistically linked to many species (e.g. oil spill and acoustic stress). Second, reducing impacts from common stressors will likely reduce risk to similar species with similar vulnerabilities. Alternatively, reducing impacts from diverse stressors will likely reduce risk to a greater diversity, and greater number, of species in an ecosystem. Species diversity also predictably contributes to the provision of ecosystem service diversity through trait diversity (23). The logic that reducing multiple risks supports higher diversity is supported by our findings: scenarios that reduce risk from diverse stressors reduce risk for more species and for the supply of more ecosystem services.

Our results indicating the predominance of leverage nodes may be generalizable despite the diversity of impact interactions. Though the risk assessment employed in our analysis uses an additive model of cumulative impacts, multiple impacts on species can interact additively, antagonistically or synergistically (24, 25). Potentially, effective management of species will require managing most stressors, such as when antagonisms persist and stressors contribute similarly to impact (24). There are also possible situations where reducing impact from any individual stressors will provide favourable management results beyond what an additive model may predict, such as when synergisms are present (26). However, a growing literature highlights the role of asymmetric and dominant stressors in regulating impact to species (27–30). Leverage nodes will occur in additive interactions where dominant stressors occur, which would be predicted directly from additive models (24, 25). Where antagonisms are found, a dominant stressor can drive the magnitude impact (27). Even in contexts of synergisms, asymmetries in impact contribution can point to dominant stressors (31).

Our study links human activities to ecosystem service provision. We are cautious to address risk to ecosystem services fully, because we only capture the biophysical aspect of ecosystem services (32). Services also have social dimensions (such as human access to services) that are prone to risks independent from biophysical risks (33, 34). We did not address these social dimensions because of the lack of available data. We used mechanistic pathways of risk through biophysical pathways, but so long as social impacts can be described through pathways, theoretically they can be represented in networks.

As in all BBN approaches, our results are dependent on the network structure we supplied. Other network structures (such as differences in links between nodes or in which other nodes are included) might change the results, so confidence in the results depends on confidence in the input network and data. The risk assessment scoring and network connections used here were created as part of a pilot project and were intended as illustrative for the potential of risk assessment and not intended for use directly by managers without further vetting. Future analyses can improve on our effort by deriving probabilities for each node that explicitly represent the possibilities for a node to change under management. Our approach employs BBNs to derive conditional probabilities of various linked states from probabilities of each node *under current management*, which offers insights but may miss important opportunities for management change (35). However, our findings that leverage nodes are more dependent on the elasticity of nodes (rather than estimates of risk probabilities) indicates that the usefulness of our methods is not restricted to cases where probability data are highly vetted but rather when the breadth of risk classes that nodes can vary among are considered thoroughly. Importantly, our methods allow for a wide range of analyses, showcased in this paper: evaluating multiple scenarios across simulated time periods. This kind of analysis cannot be done with existing network approaches to study cumulative impacts (3, 9, 10).

The emerging emphasis of strategic environmental assessments (SEA) and cumulative impact assessment (CIA) in many regulatory regimes of the world are another avenue for impact networks and leverage nodes to inform management (36, 37). Proposing mitigations is one of the most important steps in the environmental assessment process (38), and discovering leverage nodes in impact networks can help determine where mitigations are most needed for environmental protection, and where mitigations provide the best outcomes to diminish impacts from development. Determining what mitigations to apply will be context-specific, but when resources are limited, prioritization can help ensure mitigation effectiveness is maximized.

Currently, cumulative impact mapping is an important tool to aid in spatial planning (31, 39, 40). We see impact networks and leverage nodes complementing impact-mapping tools by prioritizing human activities and stressors that should be excluded from some areas. Where impact mapping analysis is done to prioritize areas to zone for protection, impact networks can help determine which human activities are most important to limit within protected areas. Networks and leverage nodes can also complement structured decision processes tasked with determining management goals and targets by providing an analytical tool to help achieve those goals (3, 41).

Effective management of cumulative anthropogenic impacts on the environment will benefit from prioritization of management actions in the context of growing human pressure on the planet.

Understanding cumulative impacts as a network of mechanistic pathways and analyzing the most prominent leverage nodes provides an effective, efficient approach to prioritize management actions. Management plans with an ecosystem-based framework, that attempt to manage cumulative impacts, can be ineffective if management actions are misallocated (21, 42). Instead of assuming leverage nodes in management plans, we advocate for revealing leverage nodes via analysis. Our research provides a practical basis to aid emerging and ongoing environmental management.

## Methods

### Bayesian Belief Networks

We used directional impact networks (with no feedback loops), assigning probabilities of risk states to nodes (e.g. a node for a species faces 40% probability of low impact and 60% high impact), and related nodes through conditional probability tables (43). A conditional probability table describes the probability of a set of nodes meeting some criteria contingent on the state of other nodes (43, 44). With a BBN, therefore, we could set the impact level on a species (or set of species) at a desired level and see what activity and stressor nodes changed the most from initial probabilities to identify the leverage nodes.

### Calculating Risk

We adapted a pilot cumulative risk assessment on 17 different species and species groups on coastal British Columbia, Canada (11, 20). Briefly, this risk assessment considered direct risk and indirect risk (i.e., risk experienced by one species is carried over to another species through ecological dependencies). Direct risk pathways are represented in Figure S2 and ecological dependencies (that regulate indirect risk) are represented in Figure S1. Human activities and stressors are listed in Table S5, and species and species groups are listed in Table S6. See Supplementary Methods for more details.

We also linked the species to 15 ecosystem services they provide (Table S6). We treated risk to ecosystem service provision (*ESR*) to be a sum of the risk of the species that contribute to each ecosystem service, normalized by the number of species contributing to *ESR*, according to the formula

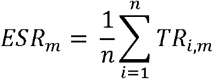

where *m* is the ecosystem service. We stress that this risk is on ecosystem service provision alone, and not the full risk to ecosystem services, as we only capture risk to the ecological side of ecosystem services, and do not account for the social dimensions (32).

Every criterion had an associated uncertainty score, and we used this score to incorporate uncertainty across the network (See Supplementary Methods) (45). The uncertainty scores are on a relative scale, and we use these scores to generate distributions of criteria. Using a Monte Carlo resampling procedure, we randomly selected values of *Temporal Scale, Spatial Scale, Intensity*, and *Consequence* from their distributions to generate a distribution of *Total Risk* for every species and ecosystem service (45). Because we knew the structure of each impact pathway and the risk from each pathway, we could also calculate how much risk originates in each activity (*OR*), and how much risk flows through each stressor (*RF*), according to the formula

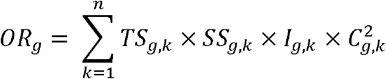

and

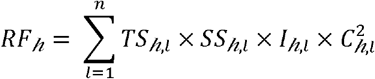

Where *g* is the human activity, *h* is the stressor, *k* is the mechanistic pathway from stressor to species, and *l* is the intersection of activity and species through stressor *h*.

### Creating the Conditional Probability Table

Using the 33^rd^ and 66^th^ percentile of risk scores within each level (activity, stressor, species, ecosystem service), we set categories of low, medium, and high risk. For example, if for a particular iteration of the Monte Carlo procedure the stressor “oil spill” has a risk score that falls above the 66^th^ percentile of the risk scores in the stressor category, then “oil spill” would be categorized as having high relative risk for that iteration. If, for the next iteration, the risk score for “oil spill” fell between the 33^rd^ and 66^th^ percentile of the risk scores in the stressor category, then “oil spill” would be categorized as having medium relative risk. Using a Monte Carlo process to generate these relative risk states among all nodes concurrently allowed us to create conditional probability tables to populate a BBN. We use these categories and cutoffs for illustrative purposes only, as we recognize that they are arbitrary designations and thresholds; however, they are useful in examining changes to risk probabilities broadly and according to the same criteria. Setting the risk levels this way meant that in some cases a node invariably faces risk at a certain level (e.g. 100% probability of being at high risk), preventing us from manipulating that node. For example, killer whales (*Orcinus orca*) consistently face comparatively high levels of risk based on our risk data, and finfish aquaculture invariably produces risk at high levels. BBN analysis was conducted using the R package gRain (46).

### Identifying Leverage Nodes

Leverage nodes for management should reflect what management strategies will drive outcomes from current conditions to goal states (15). With a BBN therefore, we can set the risk level of a species (or set of species) at a desired goal level and determine what associated activity and stressor nodes change most from initial probabilities (current conditions) to identify the leverage nodes. We define leverage nodes as those nodes that contribute over twice the average total change in risk probabilities from activity and stressor nodes when comparing initial conditions to conditions where target species are guaranteed to be at low risk. Once leverage nodes were identified, we could determine if they exist within different sectors (fisheries, sea-based activities, land-based activities, or long term stressors) based on the categories of the risk assessment (11).

We conducted a sensitivity analysis of our methods on the underlying data by determining leverage nodes for herring under multiple conditions. The first condition used the input data from the original pilot risk assessment. The second condition included a single iteration whereby all inelastic nodes in the original risk assessment are at a different risk level (e.g. a node fixed at “high risk” would have one iteration where it is “low risk”). The third condition included ten iterations where previously inelastic nodes were at a different risk level. The fourth condition included 100 iterations where nodes had this extended variation across risk levels. When leverage nodes for all conditions were determined, we calculated the similarity among conditions based on the Bray-Curtis dissimilarity index (47).

### Scenarios

We set the herring node as 100% probable of being at a low risk state and determined the activities and stressors whose risk state probabilities deviated the most from initial conditions (we call this the *target herring* scenario). Next, we compared the revealed leverage nodes as identified by the *target herring* scenario with an existing management plan (Fisheries and Oceans Canada’s (DFO) Integrated Fisheries Management Plan (IFMP)) (12). The goals of the strategy are to protect herring, while acknowledging concerns for Species At Risk (SAR) as designated by the Canadian Species at Risk Act (SARA), including Cassin’s Auklets, Steller sea lions, humpback whales and killer whales. Because killer whales in the risk assessment we base our analysis on are outliers in their ecological characteristics (having a single obligate feeding relationship with salmon) and in their risk level (facing very high risk levels), we do not include them in our analysis (20). This IFMP outlines implicit assumptions about leverage nodes in an impact network by indicating the activities and stressors it will manage. Specifically, it assumes that three activities (seine fisheries, gillnet fisheries, recreational fisheries), and one stressor (bycatch) are leverage nodes. To represent the *IFMP* scenario, we set the risk state for these assumed leverage nodes to be 100% low risk. The resulting changes to the herring node, SAR, and all species from initial conditions represent the consequences of this management plan. In comparison, we establish the *Herrign Leverage* scenario whereby we set the revealed leverage nodes for herring at 100% low risk. We compared the implications of managing revealed leverage nodes for herring in the *Herring Leverage* scenario with these *IFMP* assumed leverage nodes to predict how different management plans would perform. We also assessed predicted results to risk state probabilities for the provision of ecosystem services, paying particular attention to risk to the ecosystem service *wild harvest* as the *IFMP* is a fisheries plan.

Finally, we investigated an additional scenarios. This scenario explores the revealed leverage nodes for managing herring and SAR at low risk levels (the *target SAR* scenario). We explored how these scenarios compare with the herring-specific scenarios (*target herring* and *IFMP*). This scenario was compared with the *Herring Leverage* scenario to investigate management performance on single species vs multi-species management.

We also repeated analyses for the IFMP, Herring Leverage, and SAR Leverage scenarios for a second time step. The second time steps were path dependent, whereby the initial states of the second time step for the IFMP scenario (for example) assumed that management of the first time step in the IFMP scenario was successful. For IFMP risk from seine fishing, recreational fishing, gillnet fishing, and bycatch was minimized and the cumulative risk assessment was conducted for this new time step. From the results of this risk assessment, we set the risk states from the assumed leverage points in the IFMP to low risk, and evaluated the changes in risk profiles across species (similar to what we did in the first time step but with new risk assessment results). We created similarly path-dependent scenarios for the *Herring Leverage* and *SAR Leverage* scenarios for a second time step, whereby we assumed that management in the first time step was effective (so we reran the risk assessment with minimized risk from the leverage nodes) then determined new leverage nodes for each scenario and evaluated consequences across species from managing these new leverage nodes.

### Estimating the Species and Ecosystem Service Provision at Risk

Using the resulting probabilities of species and the provision of ecosystem services at each risk level from the BBN analysis, we established a random resampling procedure to select risk levels for each species and ecosystem service across 1000 iterations. Using these data on risk levels we calculated bootstrap 95% confidence intervals (using the BC_a_ method) of the mean (48).

## Acknowledgements

We would like to thank the Centre for Excellence for Environmental Decisions for providing a Visiting Fellowship to GGS, allowing for collaboration on this research.

## Literature Cited

1. Halpern BS, et al. (2008) A global map of human impact on marine ecosystems. Science 319(5865):948–952.

2. Sanderson EW, et al. (2002) The Human Footprint and the Last of the Wild: the human footprint is a global map of human influence on the land surface, which suggests that human beings are stewards of nature, whether we like it or not. BioScience 52(10):891–904.

3. Niemeijer D & de Groot RS (2008) Framing environmental indicators: moving from causal chains to causal networks. Environment, Development and Sustainability 10(1):89–106.

4. Brown CJ, Saunders MI, Possingham HP, & Richardson AJ (2014) Interactions between global and local stressors of ecosystems determine management effectiveness in cumulative impact mapping. Diversity and Distributions 20(5):538–546.

5. McLeod K & Leslie H (2009) Ecosystem-based management for the oceans (Cambridge Univ Press).

6. Elliott M, et al. (2017) “And DPSIR begat DAPSI (W) R (M)!”-A unifying framework for marine environmental management. Marine pollution bulletin 118(1-2):27–40.

7. Halpern BS, Lester SE, & McLeod KL (2010) Placing marine protected areas onto the ecosystem-based management seascape. Proceedings of the National Academy of Sciences 107(43):18312–18317.

8. Fulton EA, Smith AD, Smith DC, & Johnson P (2014) An integrated approach is needed for ecosystem based fisheries management: insights from ecosystem-level management strategy evaluation. PLoS ONE 9(1):e84242.

9. Knights AM, Koss RS, & Robinson LA (2013) Identifying common pressure pathways from a complex network of human activities to support ecosystem-based management. Ecological applications 23(4):755–765.

10. Cook GS, Fletcher PJ, & Kelble CR (2014) Towards marine ecosystem based management in South Florida: investigating the connections among ecosystem pressures, states, and services in a complex coastal system. Ecological Indicators 44:26–39.

11. Murray CC, Mach ME, & O M (2016) Pilot ecosystem risk assessment to assess cumulative risk to species in the Pacific North Coast Integrated Management Area (PNCIMA) (DFO Can. Sci. Advis. Sec. Res. Doc.), (DFO).

12. Kanno R (2015) Herring Integrated Fisheries Mangeement Plan. (DFO).

13. Simmonds MP, et al. (2014) Marine noise pollution-increasing recognition but need for more practical action.

14. Fagan WF & Holmes E (2006) Quantifying the extinction vortex. Ecology Letters 9(1):51–60.

15. Arkema KK, Abramson SC, & Dewsbury BM (2006) Marine ecosystem-based management: from characterization to implementation. Frontiers in Ecology and the Environment 4(10):525–532.

16. Hose JE, et al. (1996) Sublethal effects of the (Exxon Valdez) oil spill on herring embryos and larvae: morphological, cytogenetic, and histopathological assessments, 1989 1991. Canadian Journal of Fisheries and Aquatic Sciences 53(10):2355–2365.

17. Gerlotto F & Fréon P (1992) Some elements on vertical avoidance of fish schools to a vessel during acoustic surveys. Fisheries Research 14(4):251–259.

18. Slovic P (1987) Perception of risk. Science 236(4799):280–285.

19. Linkov I, et al. (2006) From comparative risk assessment to multi-criteria decision analysis and adaptive management: Recent developments and applications. Environment International 32(8):1072–1093.

20. Murray CC, et al. (2016) Supporting Risk Assessment: Accounting for indirect risk to ecosystem components. PLoS ONE 11(9):e0162932.

21. Pikitch EK, et al. (2004) Ecosystem-Based Fishery Management. Science 305(5682):346–347.

22. Zhou S, et al. (2010) Ecosystem-based fisheries management requires a change to the selective fishing philosophy. Proceedings of the National Academy of Sciences 107(21):9485–9489.

23. Cadotte MW, Cavender-Bares J, Tilman D, & Oakley TH (2009) Using phylogenetic, functional and trait diversity to understand patterns of plant community productivity. PloS one 4(5):e5695.

24. Crain CM, Kroeker K, & Halpern BS (2008) Interactive and cumulative effects of multiple human stressors in marine systems. (Wiley-Blackwell), pp 1304–1315.

25. Darling ES & Côté IM (2008) Quantifying the evidence for ecological synergies. Ecology Letters 11(12):1278–1286.

26. Crain CM, Halpern BS, Beck MW, & Kappel CV (2009) Understanding and managing human threats to the coastal marine environment. Annals of the New York Academy of Sciences 1162(1):39–62.

27. Darling ES, McClanahan TR, & Côté IM (2010) Combined effects of two stressors on Kenyan coral reefs are additive or antagonistic, not synergistic. Conservation Letters 3(2):122–130.

28. Ban SS, Graham NA, & Connolly SR (2014) Evidence for multiple stressor interactions and effects on coral reefs. Global Change Biology 20(3):681–697.

29. Folt C, Chen C, Moore M, & Burnaford J (1999) Synergism and antagonism among multiple stressors. Limnology and Oceanography 44(3):864–877.

30. Brown CJ, Saunders MI, Possingham HP, & Richardson AJ (2013) Managing for Interactions between Local and Global Stressors of Ecosystems. PLoS ONE 8(6):e65765.

31. Halpern BS, McLeod KL, Rosenberg AA, & Crowder LB (2008) Managing for cumulative impacts in ecosystem-based management through ocean zoning. Ocean & Coastal Management 51(3):203–211.

32. Tallis H, et al. (2012) New metrics for managing and sustaining the ocean’s bounty. Marine Policy 36(1):303–306.

33. Chan KM, et al. (2012) Where are cultural and social in ecosystem services? A framework for constructive engagement. BioScience 62(8):744–756.

34. Chan KM, Satterfield T, & Goldstein J (2012) Rethinking ecosystem services to better address and navigate cultural values. Ecological Economics 74:8–18.

35. McCann RK, Marcot BG, & Ellis R (2006) Bayesian belief networks: applications in ecology and natural resource management. Canadian Journal of Forest Research 36(12):3053–3062.

36. Bina O (2007) A critical review of the dominant lines of argumentation on the need for strategic environmental assessment. Environmental Impact Assessment Review 27(7):585–606.

37. Fischer TB (2010) The theory and practice of strategic environmental assessment: towards a more systematic approach (Routledge).

38. Wood C (2003) Environmental impact assessment: a comparative review (Pearson Education) p 430 pp.

39. Halpern BS, et al. (2012) Near-term priorities for the science, policy and practice of Coastal and Marine Spatial Planning (CMSP). Marine Policy 36(1):198–205.

40. Halpern BS, et al. (2009) Mapping cumulative human impacts to California Current marine ecosystems. Conservation Letters 2(3):138–148.

41. Gregory R, et al. (2012) Structured decision making: a practical guide to environmental management choices (John Wiley & Sons).

42. Game ET, Kareiva P, & Possingham HP (2013) Six Common Mistakes in Conservation Priority Setting. Conservation Biology 27(3):480–485.

43. Marcot BG, Holthausen RS, Raphael MG, Rowland MM, & Wisdom MJ (2001) Using Bayesian belief networks to evaluate fish and wildlife population viability under land management alternatives from an environmental impact statement. Forest ecology and management 153(1):29–42.

44. Jensen FV (1996) An introduction to Bayesian networks (UCL press London).

45. DFO (2015) Application of an ecological risk assessment framework to inform ecosystem-based management for SGaan Kinghlas-Bowie Seamount and Endeavour Hydrothermal Vents Marine Protected Areas. (DFO Can. Sci. Advis. Sec. Sci. Advis).

46. Højsgaard S (2012) Bayesian networks in R with the gRain package. Journal of Statistical Software.

47. Bray JR & Curtis JT (1957) An ordination of the upland forest communities of southern Wisconsin. Ecological monographs 27(4):325–349.

48. Shiue W-k, Xu C-w, & Rea BC (1993) Bootstrap confidence intervals for simulation outputs. Journal of statistical computation and simulation 45(3-4):249–255.

