## Supplementary Methods for "Prioritizing Management for Cumulative Impacts"

To explore impact networks and determine leverage nodes, we employed the following steps. First, we created an impact network based on a risk assessment for coastal BC (Murray et al. 2016a, Murray et al. 2016b). Next, we used the risk assessment to calculate conditional probability tables for a Bayesian Belief Network analysis (BBN). We then used the BBN to identify leverage nodes for the species in the network, and scenarios of multiple species. We compared the revealed leverage nodes for herring against the assumed leverage nodes in the herring *IFMP*. Finally, we assessed the consequences across the biological community and supply of ecosystem services from the various scenarios we assessed.

#### Risk Assessment

After the species were selected, all relevant stressors were listed and explicitly connected to the species based on literature reviews. The human activities driving these stressors were then linked to the stressors to create a network of pathways of effects on these species (a network of human activities →stressors→species). Risks to species come from fisheries, other ocean activities (e.g., shipping, aquaculture), terrestrial activities (e.g., land use), and long term stressors (e.g,. climate change).

After the network was assembled, a cumulative risk assessment was undertaken to quantify risk from all pathways to the species. Risk scores were literature derived, providing scores and measurements of uncertainty for each pathway. The risk calculation was made up of characteristics of exposure and consequences of impacts to species, with each score having an associated uncertainty measure.

In this risk assessment, total risk (*TR*) to each species *i* from each impact pathway *j* is a product of the criteria *Temporal Scale* (*TS*) and *Spatial Scale* (*SS*) of a stressor from an activity*, Intensity* (*I*) of the stressor from an activity*,* and *Consequence* (*C*) of the impact pathway according to the formula

$${TR}_{i}= \sum_{j=1}^{n} {TS}_{i,j}\times{SS}_{i,j}\times I_{i,j}\times C_{i,j}^{2}$$

Because exposure criteria (*TS, SS,* and *I*) are on the scales 0-4, 0-3, and 0-3, respectively, and the consequence criterion (*C*) is on a 0-6 scale, *C* is squared to give exposure and consequence equal weighting (Murray et al. 2016a, Murray et al. 2016b). To incorporate indirect risk, we assumed that risk flowed through trophic links of species according to 10% rule, so that predators accumulated 10% risk of prey (Lindeman 1942). We also assumed that habitat-providers transfer 10% of their risk to habitat-receptors (Figure S1).

Though this risk assessment provided us with the best available data to explore impact networks and leverage nodes, this risk assessment was originally intended to characterize current risk to species, with uncertainty measures characterizing certainty in particular measures of risk. We use the final risk distribution derived from these scores of risk criteria and associated uncertainty as a proxy for the scope of change in risk throughout the network, which is necessarily smaller than the actual scope of change for risk. As a result, our findings should not be taken as conclusive for specific management but rather illustrative of the application of our methods.

#### Scoring exposure and consequence

The data on risk scores for this project were based on a risk assessment conducted for coastal British Columbia (Clarke Murray et al. 2016). Below, we outline the prominent components of risk, their qualitative values, and calculating final risk scores from these criteria (with associated uncertainty) using a Ecological Risk Assessment Framework developed by O et al (2015) and adapted by Clarke Murray et al (2016).

*Temporal Scale (TS)* refers to the frequency of the event, rather than its duration. Consideration was given to how often the stressor occurs, rather than how long the effect is felt by the SEC (which in practice was included in the *Consequence_ij_* scoring). Scoring is described in Table S1A.

*Spatial Scale* *(SS)* is the scale or spatial extent of the impact from the stressor. For example, under sedimentation from trawl fisheries, consideration was given to how far sediment is carried from the site of the trawl. Scoring for the dive fishery considered the size of the footprint of habitat disturbance from a single dive. Scoring is described in Table S1B.

*Intensity (I)* is a measure of the density and persistence of the stressor. Depending on the stressor or activity in question, Intensity can refer effort, density, amount of an activity, or the amount or strength of a stressor (e.g. quantity or concentration of a pollutant or harmful species, rate of change for climate change) across the entire study area. For example, load for finfish aquaculture evaluates how many finfish farms there are in British Columbia and how often and how much area is covered by finfish farms. Scoring described in Table S1C.

*Consequence* (*C*) is the impact of the stressor on the individual environmental component and therefore must be scored for each environmental component by each mechanistic impact pathway. It is scored from 1 to 6 ranging from negligible to intolerable consequence and indicates the impact of the stressor on the individual SEC, as described in Table S1D. *Consequence_ij_* scoring is based on the subcomponent (population size, geographic range, behaviour, etc) but most commonly *Consequence_ij_* was scored on the population size or geographic range subcomponent. If information was available about more than a single subcomponent, the most sensitive subcomponent was used to assign the score. In choosing the most sensitive subcomponent, consideration should be given to the subcomponent most important for long-term persistence and/or the subcomponent that is the most sensitive to the stressor being scored. Uncertainty was also included for the *Consequence_ij_* score; see Table S2 for uncertainty categories and scores.

Table S1. Scoring of variables: a) Spatial Scale, b) Temporal Scale, c) Load and d) *Consequence*

| **(a) *Temporal Frequency Scale*** | | | |
| --- | --- | --- | --- |
| **Score** | **Effect** | | **Definition** |
| 1 | Rare | | Every several years – Decadal |
| 2 | Relatively Often | | Quarterly – Annually |
| 3 | Frequent | | Weekly – Monthly |
| 4 | Continuous | | Daily occurrences or continuous |
| **(b) *Spatial Scale*** | | | |
| **Score** | **Effect** | | **Definition** |
| 1 | Few restricted locations | | 1-10 kilometres |
| 2 | Localized | | 10-100 kilometres |
| 3 | Widespread | | >100 kilometres |
| **(c) *Load – Density/Persistence*** | | | |
| **Score** | **Effect** | | **Definition** |
| 1 | Low | | Low density and low persistence |
| 2 | Moderate | | High density or persistence |
| 3 | High | | High density and persistence |
| **(d) *Consequence*** | | | |
| **Score** | **Effect** | **Definition** | |
| 1 | Negligible | Negligible impact on population/habitat/community | |
| 2 | Minor | Minimal impact on population/habitat/ community structure or dynamics | |
| 3 | Moderate | Maximum impact that still meets an objective (e.g. sustainable level of impact such as a full exploitation rate for a target species; maintaining levels of critical habitat) | |
| 4 | Major | Wider and longer term impacts (e.g. long-term decline in CPUE) | |
| 5 | Severe | Very serious impacts occurring, with a relatively long time period likely to be needed to restore to an acceptable level (e.g. serious decline in spawning biomass limiting population increase) | |
| 6 | Intolerable | Widespread and permanent/irreversible damage or loss will occur – unlikely to ever be fixed (e.g. local extinction) | |

Table S2. Scoring definition of the uncertainty of risk scores.

| ***Uncertainty*** | | |
| --- | --- | --- |
| **Score** | **Literature** | **Definition** |
| 1 | Extensive | Extensive scientific information; peer-reviewed information; data specific to the location; supported by long-term datasets (10 years or more) |
| 2 | Substantial | Substantial scientific information; non-peer-reviewed information; data specific to the region; supported by recent data (within the last 10 years) or research |
| 3 | Moderate | Moderate level of information; data from comparable regions or older data (more than 10 years) from the area of interest |
| 4 | Limited | Limited information; expert opinion based on observational information or circumstantial evidence |
| 5 | Little to None | Little or no information; expert opinion based on general knowledge |

#### Scoring Uncertainty

An uncertainty incorporation exercise was completed to include the uncertainty of the qualitative risk scoring in the final and cumulative risk scores. Each risk variable (*Temporal Scale, Spatial Scale, Intensity* and *Consequence*) was assigned as the mean of a normal distribution with standard deviation set according to the level of uncertainty assigned (Table S2). The distribution was bounded by the minimum and maximum scores for each risk variable so that the scores could not be higher or lower than the variable’s scale (*e.g.*, the intensity score cannot be lower than 1 or higher than 3). The score of each risk variable was then randomly sampled from this distribution with 3000 replicates. The final risk score for each mechanistic impact pathway was a product of the four risk variable arrays (*Risk* = *SS* x *TS* x *I* x *C*^2^), where the first score generated from each variable array is multiplied across all four risk variables, followed by the second, and so on for all 3000 replicates and resulting in a final risk array of 3000 scores.

### Supplementary Figures and Tables

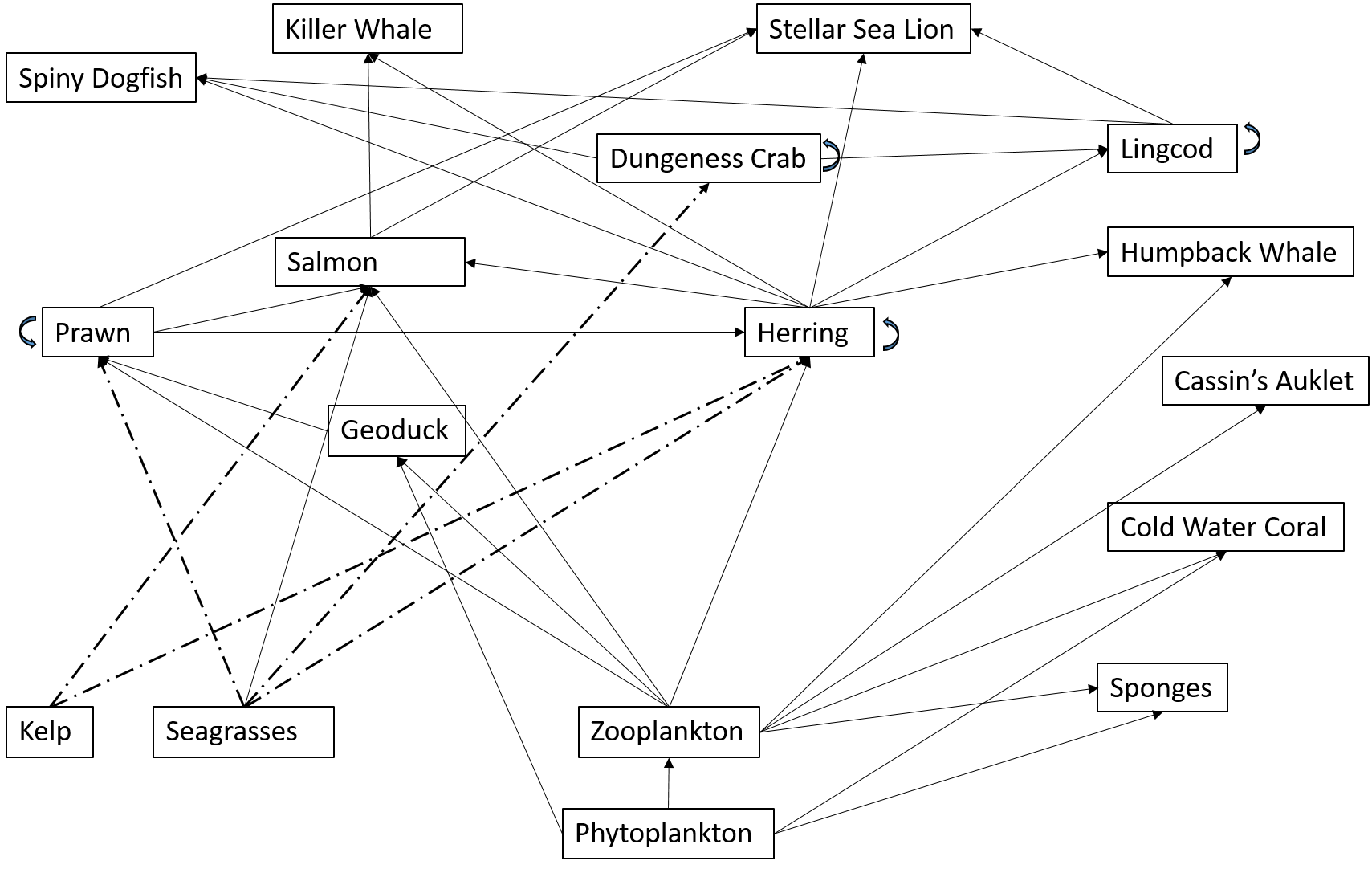

Figure S1. Food web and habitat relationships between the species in the coastal British Columbia case study. Habitat relationships are represented by dashed lines and trophic relationships are represented by solid lines (Figure adapted from Clarke Murray et al 2016).

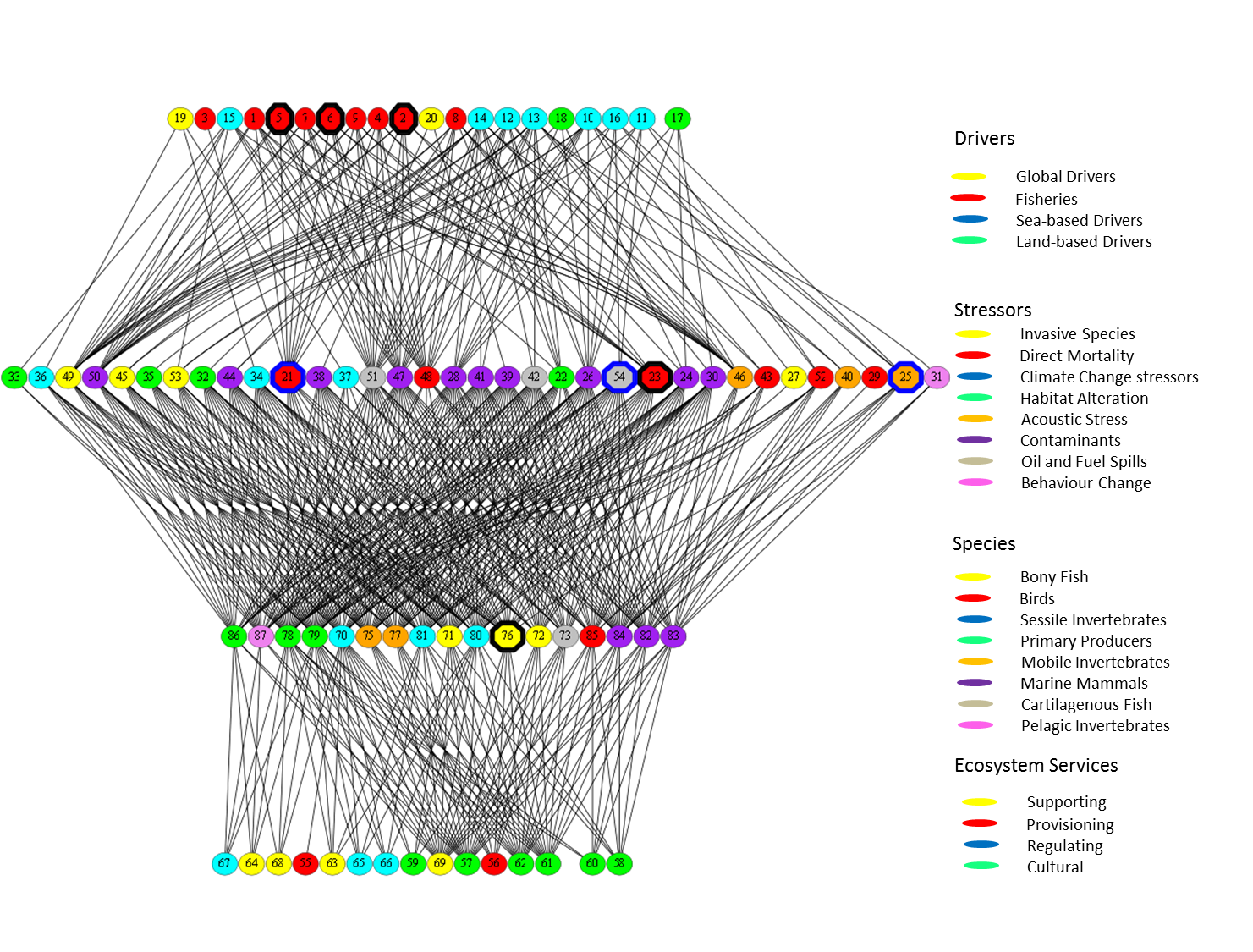

Figure S2. The impact network for 87 linking drivers, stressors, species and ecosystem services of the categories listed on the right (a given category may have multiple items, e.g., the driver “fisheries” includes seine fisheries, gillnet fisheries, and others). Numbers in the nodes correspond to the codes on Supplementary Tables S2 and S3. Nodes focused on by the Herring Integrated Fisheries Management Plan are outlined in black, while the top three revealed leverage nodes are outlined in blue.

Table S3. The leverage nodes for species, grouped by sector (fisheries, sea, land, and long term impacts). The number in brackets indicates the number of leverage nodes revealed through our analysis for each species.

| Species (number of leverage nodes) | Fisheries | Sea | Land | Long Term |
| --- | --- | --- | --- | --- |
| Herring Leverage (7) | Direct Capture | Acoustic stress; Oil spill; Change in water flow; Nutrient input | Nutrient input | Marine debris; Temperature change |
| Geoduck Clam (8) | Habitat disturbance; Sedimentation; Direct capture | Large vessel invasive species; Habitat disturbance; Invasive species; Nutrient input | Change in freshwater flow; Sedimentation; Nutrient Input | Marine debris |
| Cassin's Auklet (2) | Small vessel acoustic | Small vessel acoustic |  | Marine debris |
| Cold Water Coral (10) | Sedimentation; Small vessel incidental mortality | Large vessel incidental mortality; Incidental mortality; Small vessel incidental mortality; Nutrient input; Oil spill; Invasive species; Change in water flow | Sedimentation; Nutrient input | Marine debris; Temperature change |
| Humpback Whale (1) | Small vessel acoustic | Small vessel acoustic |  |  |
| Kelp (9) | Direct Capture; Sedimentation; Small vessel incidental mortality; Habitat disturbance | Invasive species; Small vessel incidental mortality; Habitat disturbance; Oil spill; Large vessel incidental mortality | Sedimentation | Marine debris; Sea level rise |
| Prawn (9) | Direct capture; Sedimentation; Seine Fisheries | Nutrient input; Change in water flow; Oil spill | Nutrient input; Sedimentation; Change in freshwater flow | Temperature change; Marine debris |
| Steller Sea Lion (3) | Small vessel acoustic | Small vessel acoustic; Large vessel incidental mortality; Acoustic stress |  |  |
| Seagrasses (11) | Direct capture; Small vessel incidental mortality; Sedimentation | Large vessel incidental mortality; small vessel incidental mortality; Nutrient input; Incidental mortality; Oil spill; Change in water flow | Nutrient input; Sedimentation | Marine debris; Temperature change; Sea level rise |
| Sponges (9) | Small vessel incidental mortality; Sedimentation | Large vessel incidental mortality; Incidental mortality; Nutrient input; Oil spill; Small vessel incidental mortality; Change in water flow | Nutrient input; Sedimentation | Marine debris; Temperature change |
| Zooplankton (7) |  | Oil spill; Change in water flow; Nutrient input | Nutrient input | Temperature change; Marine debris; Ocean acidification; Persistant organic pollutants |
| SAR Leverage scenario (5) | Direct capture; Small vessel acoustic | Small vessel acoustic; Acoustic stress; Oil spill; Nutrient input | Nutrient input |  |
| T2 Herring Leverage (8) | Direct capture | Nutrient input; Oil spill | Nutrient input; Sediment | Debris; Temperature change; Persistent organic pollutants |
| T2 SAR Leverage (11) | Direct capture; Habitat disturbance | Oil spill; Acoustic stress; Large vessel contaminants; Large vessel incidental mortality | Debris; Sediment | Persistent organic pollutants; Temperature change; Marine debris |

Table S4. Nodes ranked by influence in maintaining low risk to herring. The columns identify the probabilities of low, medium, and high risk according to each scenario. Nodes in bold are assumed leverage nodes in the herring IFMP.
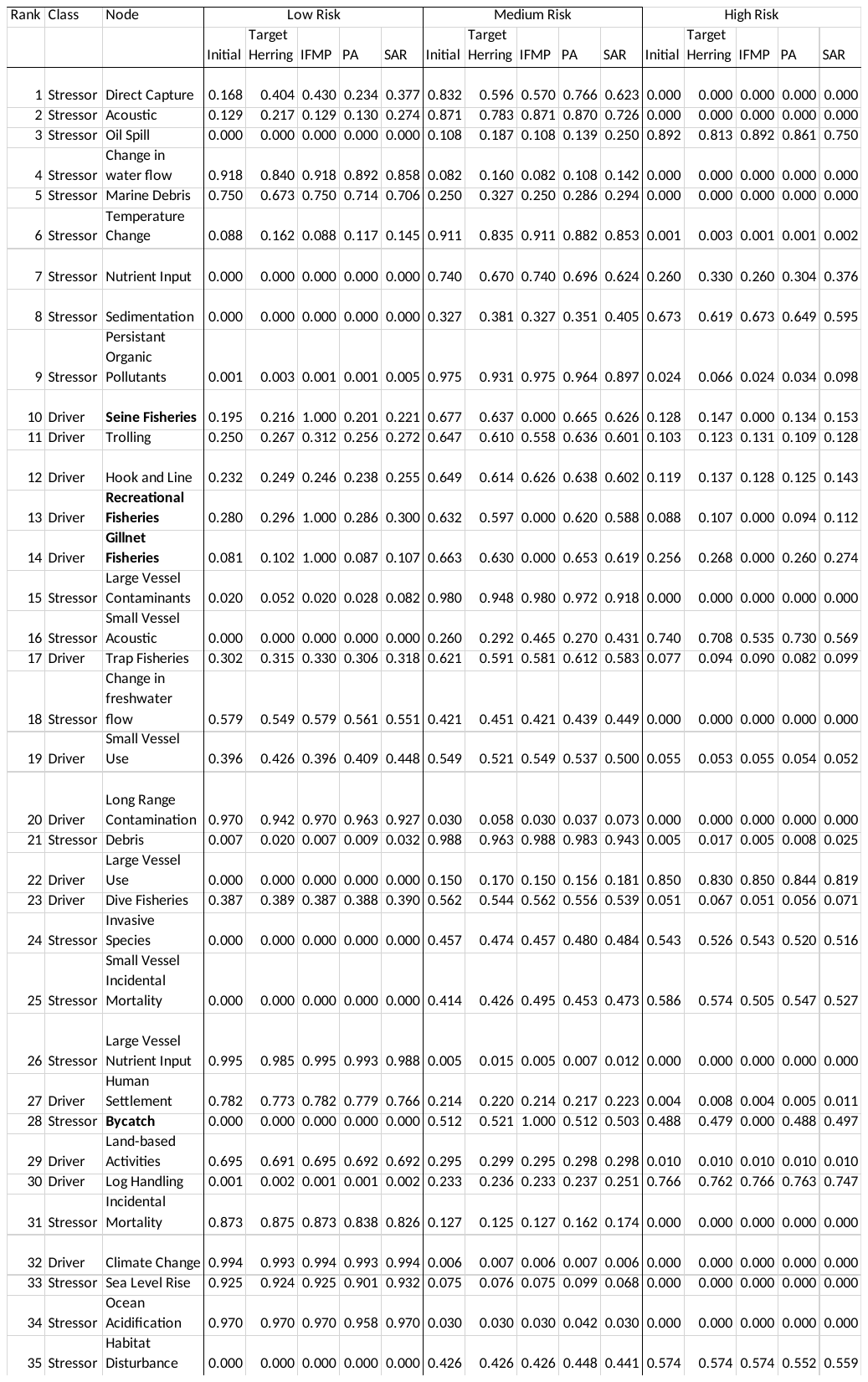

Table S5. Human activities (in bold) and associated stressors. The numbers beside each activity and stressor corresponds to the nodes in Figure 2

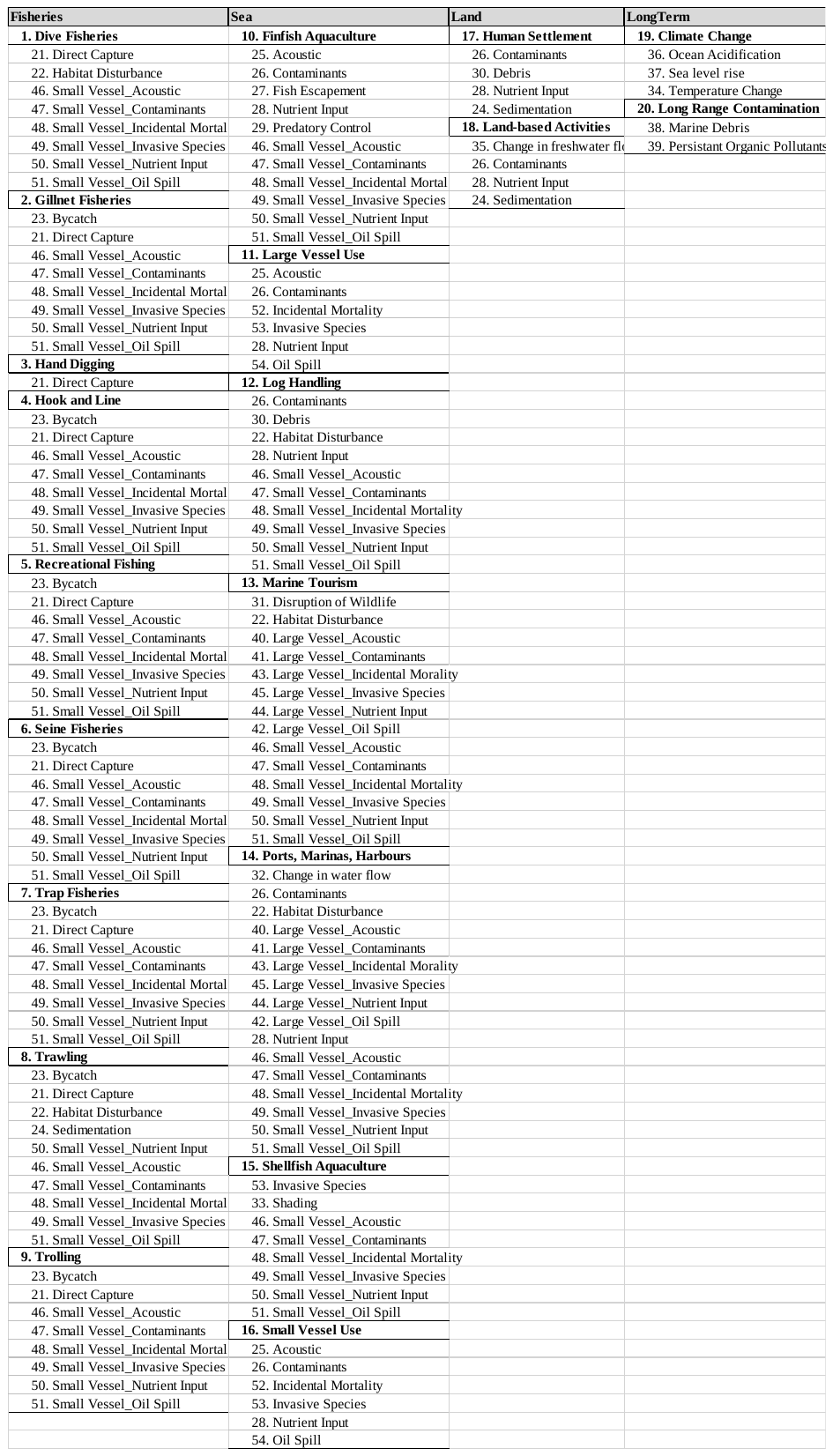

Table S6. The provision of ecosystem services (in bold) and their associated species. The numbers beside each ecosystem service and species corresponds to the nodes in Figure 2.

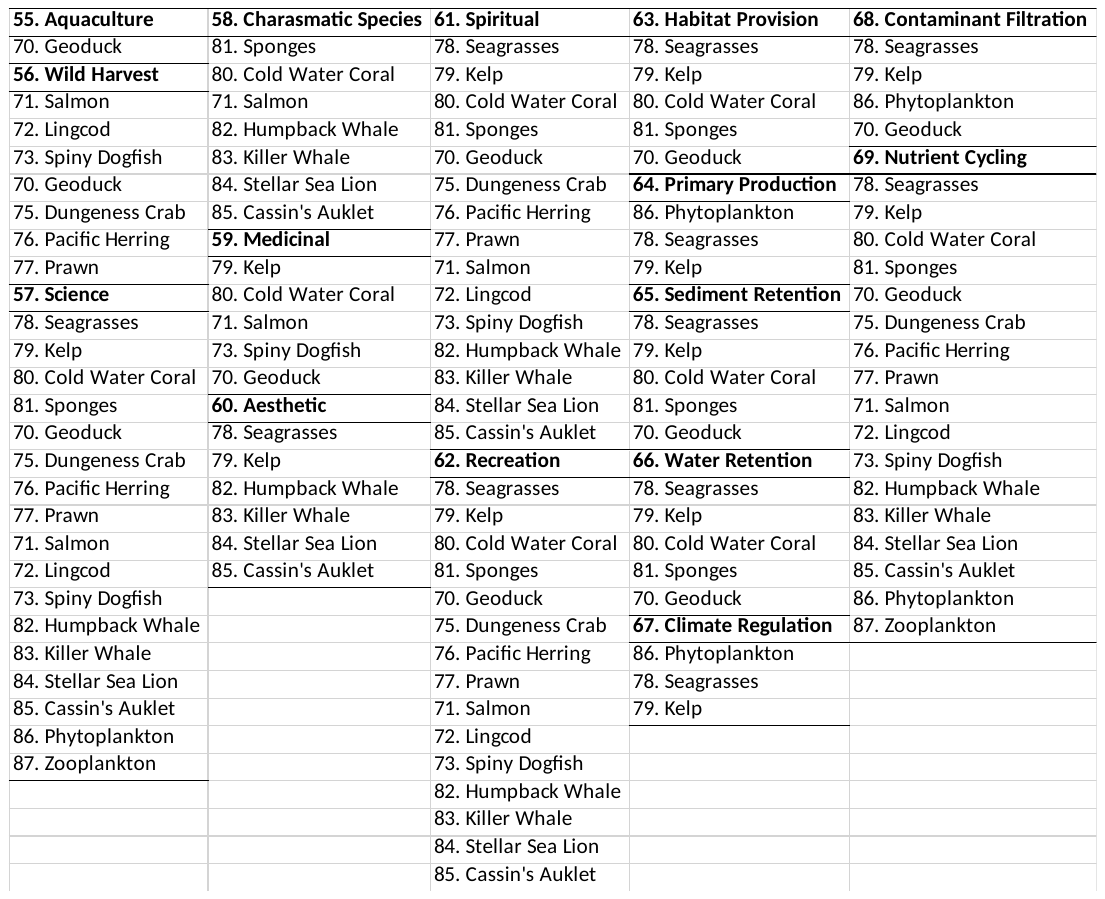

Table S7. Similarity of leverage nodes for herring among risk assessments, including the original assessment, an assessment with 1 iteration that expands the range of risk levels for nodes that are originally fixed, an assessment with 10 iterations expanding node ranges, and an assessment with 100 iterations expanding node ranges. Bray-Curtis dissimilarity values are presented, where a 0 represents no difference in leverage nodes among assessments, and 1 represents no overlap in the leverage nodes.

|  | Initial assessment | Initial + 1 iteration | Initial + 10 iterations |
| --- | --- | --- | --- |
| Initial + 1 iteration | 1 |  |  |
| Initial + 10 iterations | 1 | 0.08929742 |  |
| Initial + 100 iterations | 1 | 0.2241811 | 0.13763905 |

Table S8. Leverage nodes for herring among risk assessments, including the original assessment and assessments with iterations that expand the range of risk levels for nodes that are originally fixed. Nodes are ranked by their level of influence on risk to herring.

| Rank | Initial | Initial + 1 iteration | Initial + 10 iterations | Initial + 100 iterations |
| --- | --- | --- | --- | --- |
| 1 | Direct Capture | Small vessel contaminants | Small vessel contaminants | Small vessel contaminants |
| 2 | Acoustic | Small vessel nutrient input | Small vessel nutrient input | Small vessel nutrient input |
| 3 | Oil Spill | Small vessel oil spill | Small vessel oil spill | Small vessel oil spill |
| 4 | Change in water flow | Gillnet fisheries | Seine fisheries |  |
| 5 | Marine debris | Seine fisheries |  |  |
| 6 | Temperature change |  |  |  |
| 7 | Nutrient input |  |  |  |

#### Literature Cited

Lindeman, R. L. 1942. The trophic-dynamic aspect of ecology. Ecology **23**:399-417.

Murray, C. C., M. E. Mach, R. G. Martone, G. G. Singh, M. O, and K. M. A. Chan. 2016a. Supporting Risk Assessment: Accounting for indirect risk to ecosystem components. PLoS ONE **11**:e0162932.

Murray, C. C., M. E. Mach, and M. O. 2016b. Pilot ecosystem risk assessment to assess cumulative risk to species in the Pacific North Coast Integrated Management Area (PNCIMA). Page vii + 61 p. *in* DFO, editor. DFO Can. Sci. Advis. Sec. Res. Doc.
